# GABA_A_ receptors are selectively expressed in NG2 glia of the cerebellar white matter

**DOI:** 10.1101/741140

**Authors:** Labrada-Moncada Emmanuel, Martínez-Torres Ataulfo, Reyes-Haro Daniel

**Author notes:** For correspondence: Dr. Daniel Reyes Haro. Departamento de Neurobiología Celular y Molecular. Instituto de Neurobiología, Universidad Nacional Autónoma de México, Campus Juriquilla, Boulevard Juriquilla 3001, Juriquilla, Querétaro CP76230, México. Declaration of interest: none.

## Abstract

The cerebellum is involved in the coordination of movement. Its cellular composition is dominated by GABAergic neuronal types, and glial cells are known to express functional receptors. GABAergic signaling regulates cell proliferation, differentiation, and migration during neurodevelopment. However, little is known about the functional expression of GABA receptors in the cerebellar white matter (WM). Thus, the aim of this study was to test whether glial cells express functional GABA receptors during postnatal development (P7-P9) of cerebellar WM. Immunofluorescence studies showed that half of the astrocytes express GAD67, suggesting that GABA is synthetized by glial cells. Calcium imaging in cerebellar slices revealed that GABA and the GABA_A_ agonist muscimol evoked calcium transients in sulforhodamine B (SRB) negative cells, whereas the GABA_B_ agonist baclofen failed to evoke responses in cerebellar WM. Whole-cell patch-clamp recordings of GFAP^+^ cells showed dye coupling and a passive current-voltage relation typical of astrocytes. Surprisingly, these cells did not respond to muscimol. Two additional populations were identified as GFAP^−^ cells. The first population showed dye coupling, slow decaying inward and outward currents with no voltage dependence and did not respond to GABA_A_ agonists. The second population showed an outward-rectifying current-voltage relationship and responded to muscimol, but dye coupling was absent. These cells received synaptic input and were NG2^+^, but evoked calcium waves failed to modulate the frequency of sPSCs or signal directly to NG2 glia. We conclude that GABA_A_ receptor-mediated signaling is selective for NG2 glia in the WM of the cerebellum.

## Introduction

The cerebellum is responsible for controlling movement and maintaining balance by integrating sensory information from different regions of the central and peripheral nervous system (CNS and PNS). The cellular organization of the cerebellum includes granular and unipolar neurons that synthesize and release glutamate, while the Purkinje, Stellate, Basket, Lugaro and Golgi neurons synthesize and release gamma-aminobutyric acid (GABA) (Takayama & Inoue, 2005).

In the white matter (WM) glial cells are predominant (Sturrock, 1976; Reyes-Haro et al., 2013). Here, astrocytes are known to have a neurogenic activity that is restricted from postnatal days 2 to 12 (P2-P12). During this period, astrocytes are the source of origin of the interneurons that make up the molecular layer of the cerebellum (Silbereis et al., 2009). During postnatal development (P12-P23), oligodendrocyte precursor cells (OPCs), also known as NG2 glia, differentiate into oligodendrocytes (Foran and Peterson 1992, Nishiyama et al., 2009), which are known to be a major cell population in WM (Silbereis et al 2009).

In glial cell precursors, GABA signaling through GABA_A_ receptors (GABA_A_Rs) is known to be involved in proliferation, differentiation, and migration (Kilb et al., 2013; Bolteus and Bordey, 2004). Classically, the astrocyte-like precursor cells are responsive to GABA in multiple brain regions, such as the hippocampus (Seri et al., 2001, Ge et al., 2006) and sub-ventricular zones (Bolteus and Bordey, 2004, Liu et al., 2005). Functional expression of GABA_A_Rs has been demonstrated in diverse glial cells from the cerebellum, for example in Bergmann’s glia (Müller et at., 1994), in ependymal cells from the periventricular zone (Reyes-Haro et al., 2013) and in primary cultures of cerebellar astrocytes (Pétriz et al., 2014). Moreover, the functional expression of GABA_A_Rs in NG2 glial cells from cerebellar WM was also reported (Zonouzi et al., 2015).

Considering the fundamental role of GABA during early brain development and its critical role for glial and neuronal communication, we explored whether three types of glial cells, namely, astrocytes, oligodendrocytes, and NG2 cells, express functional GABA_A_Rs in the cerebellar WM (*in situ*).

## Methods

### Acute Brain Slice Preparation

The experiments were conducted in accordance with the guidelines approved by the Institutional Animal Care and Use Committee of the Universidad Nacional Autónoma de México. Acute brain slices were prepared from P7-P9 of CD1 mice or transgenic GFAP− eGFP mice (Nolte et al., 2001; Reyes-Haro et al., 2013a). In brief, mice were decapitated and brains were carefully removed and transferred to a chamber with ice-cold oxygenated artificial cerebrospinal fluid (aCSF) containing (in mM): NaCl 134; KCl 2.5; MgCl_2_ 1.3; CaCl_2_ 2; K_2_HPO_4_ 1.25; NaHCO_3_ 26; D-glucose 10; pH 7.4. The buffer solution was oxygenated with carbogen (95% O2, 5% CO2). Coronal slices of the cerebellum (250 μm thick) were prepared at 4 °C using a vibratome (VS100, Leica, Germany). They were then gently transferred and stored in aCSF at room temperature (20–22 °C) for at least 30 min before the experiments were performed. Each recording was made from one slice. For all the experiments, “n” refers to the number of slices and “N” to the number of animals.

### Astrocyte staining with sulforhodamine

Mice were injected intraperitoneally with sulforhodamine B (SRB; 20 mg/kg, Sigma S1402) 4 h prior to the preparation of brain slices: this approach was highly efficient for astrocyte staining as previously reported (Appaix et al., 2012).

### Dye-Coupling

Acute brain slices were placed in a holding chamber mounted on the stage of an upright light microscope (Olympus BX51W). To maintain constant conditions during experiments, the chamber was continuously perfused with standard aCSF. Patch pipettes were pulled from borosilicate capillaries (inner diameter 0.86 mm; outer diameter 1.5 mm; Sutter Instrument) using a P-97 pipette puller (Sutter Instrument) and filled with pipette solution containing (in mM): NaCl 4; KCl 120; MgCl_2_ 4; CaCl_2_ 0.5; Hepes 10; EGTA 5; D-glucose 5; 0.5% biocytin (B4261, Sigma) at pH 7.4. For some experiments, 0.1% SRB was added to the pipette solution to reveal the cell morphology, but we mostly used biocytin for this purpose. Pipette resistance ranged from 5 to 8 MΩ. A single cell per slice was filled via the patch pipette during whole-cell recordings.

The membrane was de- and hyperpolarized between −140 and +60 mV in 20 mV steps for 60 ms at the beginning of the experiment to obtain the current profile of the recorded cell. Current signals were amplified (EPC10 amplifier, HEKA), filtered (2.9 kHz) and sampled (20 kHz), and for the drug response test, the currents were sampled (2 kHz) for 5 minutes and monitored with PatchMaster software (HEKA).

### Calcium imaging

The fluorescent calcium dye Fluo-4 AM (10 μM, Life Technologies, Ex 495/506 nm) was loaded directly onto the slices and maintained for 30 min in an incubator that was oxygenated continuously with 95% O_2_, 5% CO_2_ at 37 °C in a loading chamber (1.5 cm diameter; 500 μl volume of aCSF). Following dye loading, the tissue was washed with oxygenated aCSF for at least 20 min and kept there until used.

For imaging, tissue was placed in a submerged recording chamber under an Olympus BX51W microscope and perfused with oxygenated aCSF at a flow rate of 2–3 ml/min at room temperature (~25 °C). To mitigate the movement in the stream, the slices were kept under a harp-like made of nylon threads glued across a U-shaped piece of stainless steel. Image time series were acquired with water-immersion objectives at 1 Hz.

Calcium signals were elicited with GABA 500 μM (Sigma, A2129), 50 μM muscimol (Sigma, M1523), 20 μM baclofen (Sigma, G013), 100 μM ATP (Sigma, A3377) or electrical stimulation by 20 pulses of 200 μA at 10 Hz (DS3 Isolated Current Stimulator, Digitimer Ltd) with a glass electrode filled with aCSF. The stimulation pipette (tip opening ~20 μm) was gently allocated on top of the slice touching the upper cell layer. The slice was allowed to recover from mechanical stress for at least 5 min.

Acquisition protocols consisted of 300 s and 200 s for stimulated calcium elicitations, acquiring one picture every second using a LED source (X-Cite XLED1, Excelitas Technologies). All analyses and processing were made using ImageJ/FIJI software. To visualize the spatial and temporal changes in calcium resulting from the elicited activity, the raw sequences were processed to highlight changes in fluorescence intensity between frames. Regions of interest over the field of view were selected, and the mean pixel intensity at each frame was measured. The data were first plotted as fluorescence intensity versus time and subsequently converted to a relative scale (∆F/F baseline). To allow comparison across slices, the same threshold value was used for all slices.

### Immunofluorescence

After recording and dialysis, the slices were fixed for 1 to 2 h in a solution of 4% paraformaldehyde in 0.1 M phosphate buffer (PFA-PB, pH 7.4) at 4 °C and processed for immunofluorescence. After washing the slices with PB, slices were incubated in a solution containing 1% Triton X-100 (TX-100), 5% donkey serum and 10% fetal bovine serum (FBS) in PB, at pH 7.4, for 4 h to permeabilize and block non-specific binding. Finally, biocytin was labeled with DyLight 594 conjugated streptavidin (1:200, Vector Labs, SA-5594) to reveal if the recorded cell was coupled or not.

For the NG2 antibody protocol, the slices were fixed 1 h in 0.1M PFA-PB (pH 7.4), the slices were rinsed in PBS 3 times for 10 min, incubated in a solution containing 1% TX-100, 5% donkey serum and 10% FBS in phosphate buffer at pH 7.4, for 2 h at room temperature. After washing, the slices were incubated in the antibody solution (Anti-NG2 Chondroitin Sulfate Proteoglycan, Rabbit, 1:250, AB5320, Merck Millipore) for 48 h at 4 °C, washed and moved to a solution with the secondary antibody (Alexa 488 Donkey anti-rabbit, 1:500, Thermo Fisher) and DyLight 594 conjugated streptavidin (1:200, Vector Labs, SA-5594) for 36 h at 4 °C; after, the slices were washed and mounted. For the acquisition of confocal images, a Zeiss LSM 780 DUO Confocal Microscope was used with Zeiss Zen Black software (Zeiss, Göttingen, Germany) and an oil immersion 25X objective (LCI Plan-Apochromat, NA 0.8) with DyLight 594 (DPSS Laser, excitation/emission wavelength 590/617nm) and eGFP (Argon Laser, excitation/emission wavelength 488/511nm).

GAD67 immunofluorescence was carried out with a standard method for fixed sections, in brief, the P7-9 GFAP−eGFP mice were perfused with PB 0.1 M and PFA 4%, the brains were carefully extracted and transferred into a cryoprotectant solution (PB + 30% sucralose) for 3 days, sliced in a cryostat (35 μm), then, the sections were washed, incubated in a solution containing 1% TX-100, 5% donkey serum and 10% FBS in phosphate buffer at pH 7.4, for 2 h at room temperature, washed and placed in PB with the primary antibody (anti-GAD67, mouse, 1:500, MAB5406, Merk Millipore) overnight at 4 °C, after washing, the slices were incubated for 2 hours in a solution with the secondary antibody (Alexa 594, anti-mouse, 1:1000, Thermo Fisher) at room temperature, washed and mounted.

## Results

### GAD67 is highly expressed in cerebellar white matter

GABAergic transmission requires GABA synthesis and thus we tested by immunofluorescence the expression of GAD67 in glial cells. This showed high expression in WM and low expression in the cerebellar cortex of GFAP−eGFP mice (P7-P9) (fig. 1). Astrocyte density was estimated (1133 ± 137 GFAP^+^ cells/mm2), and 54% of the GFAP^+^ cells were GAD67+ (616 ± 137 GFAP^+^/GAD67+ cells/mm2; n=6; N=4), indicating that a fraction of glial cells from the WM, including astrocytes, can synthesize GABA.

**Figure 1:**
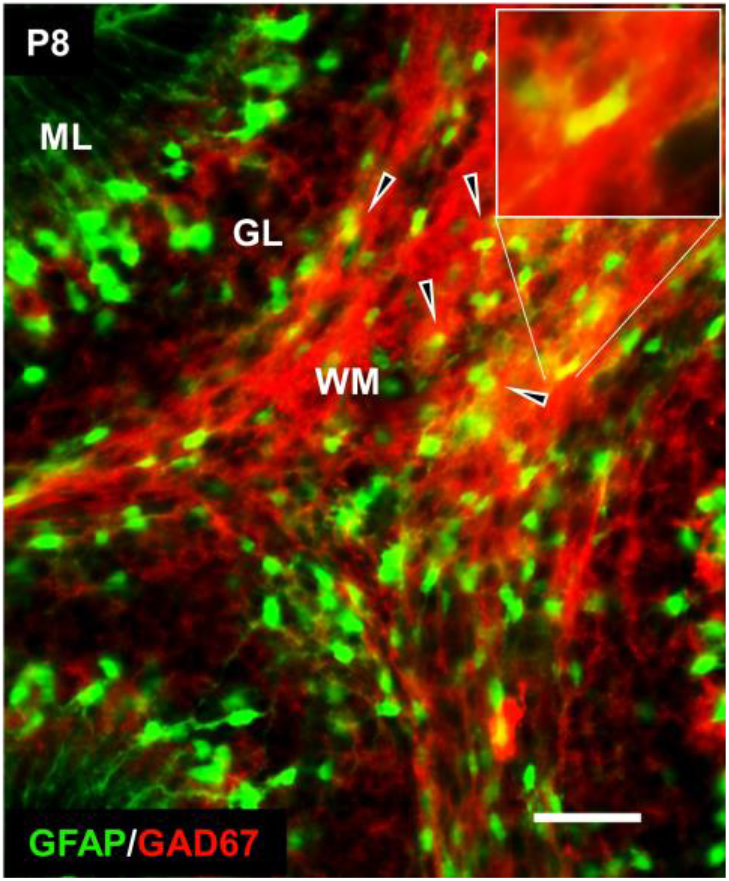
GAD67 is expressed in cerebellar WM. Immunofluorescence for GAD67 in WM of a postnatal 8 (P8) GFAP-eGFP mouse. Notice that GAD67 expression is predominant in WM and 54% of the WM astrocytes (GFAP-eGFP^+^ cells) co-localized with GAD67 expression (scale bar = 100 μm).

### GABA_A_ receptors are functionally expressed in glial cells from cerebellar white matter

The functional expression of GABA_A_Rs in glial cells from WM was tested by measuring calcium transients evoked by agonists. GABA (500 μM) added to the aCSF evoked intracellular calcium responses in 41.7% ± 5.1% of the glial cells, indicating the functional expression of GABA receptors (n=13; N=10). As a side note, the GABA-mediated response was also observed in the granular cell layer (red arrowheads, Fig. 2A). The next step was to test the agonist muscimol (50 μM) and baclofen (20 μM) for GABA_A_Rs and GABA_B_Rs respectively. The application of muscimol elicited a calcium increase in 39% ± 7% of the cells (n=10; N=6, Fig. 2B), in contrast, baclofen failed to elicit any detectable response (Fig. 2C), all these observations indicated that GABA-mediated responses are through GABA_A_Rs. ATP (100 μM) was applied at the end of the experiment to confirm slice viability, and the total number of cells that responded to ATP was considered as 100% (Fig. 2D).

**Figure 2.**
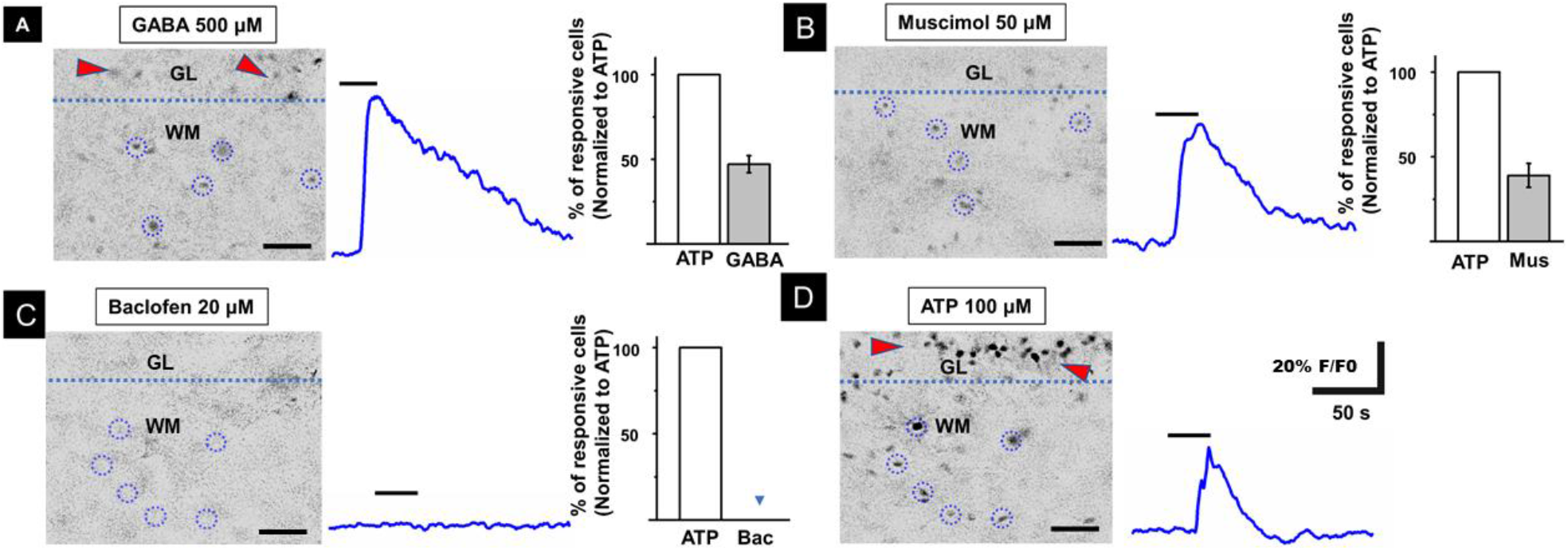
Glial cells respond to muscimol but not to baclofen. **A-** GABA (500 μM) induced a calcium response in 41.7% ± 5.1 of the WM cells (n=13; N=10). Red arrows highlight GABA-mediated responses in the granular layer (GL). **B, C**- Muscimol (50 M) evoked a calcium response in 39% ± 7% of the WM cells (n=10; N=6), while baclofen (20 M) failed to induce a response (n=10; N=6) **D-** ATP (100 M) was applied after GABAergic agonists, and the evoked response was considered as 100% of the responding WM cells. ATP also helped to verify slice viability (scale bar = 50 μm).

### Muscimol responding cells are not SRB^+^

Our next step was to test if SRB^+^ cells responded to muscimol, which would give an idea of the population of astrocytes expressing GABA_A_Rs. Unexpectedly, calcium imaging experiments showed that muscimol (50 μM) elicited responses only in SRB^−^cells, not in SRB^+^ cells (n=6; N=5). Altogether, these results suggest that astrocytes do not express GABA_A_Rs (Fig. 3A). As a positive control, we tested ATP(100 μM) at the end of the experiment and found that 33.97% ± 3.5% of the responsive cells were also SRB^+^ (n=6; N=5).

**Figure 3.**
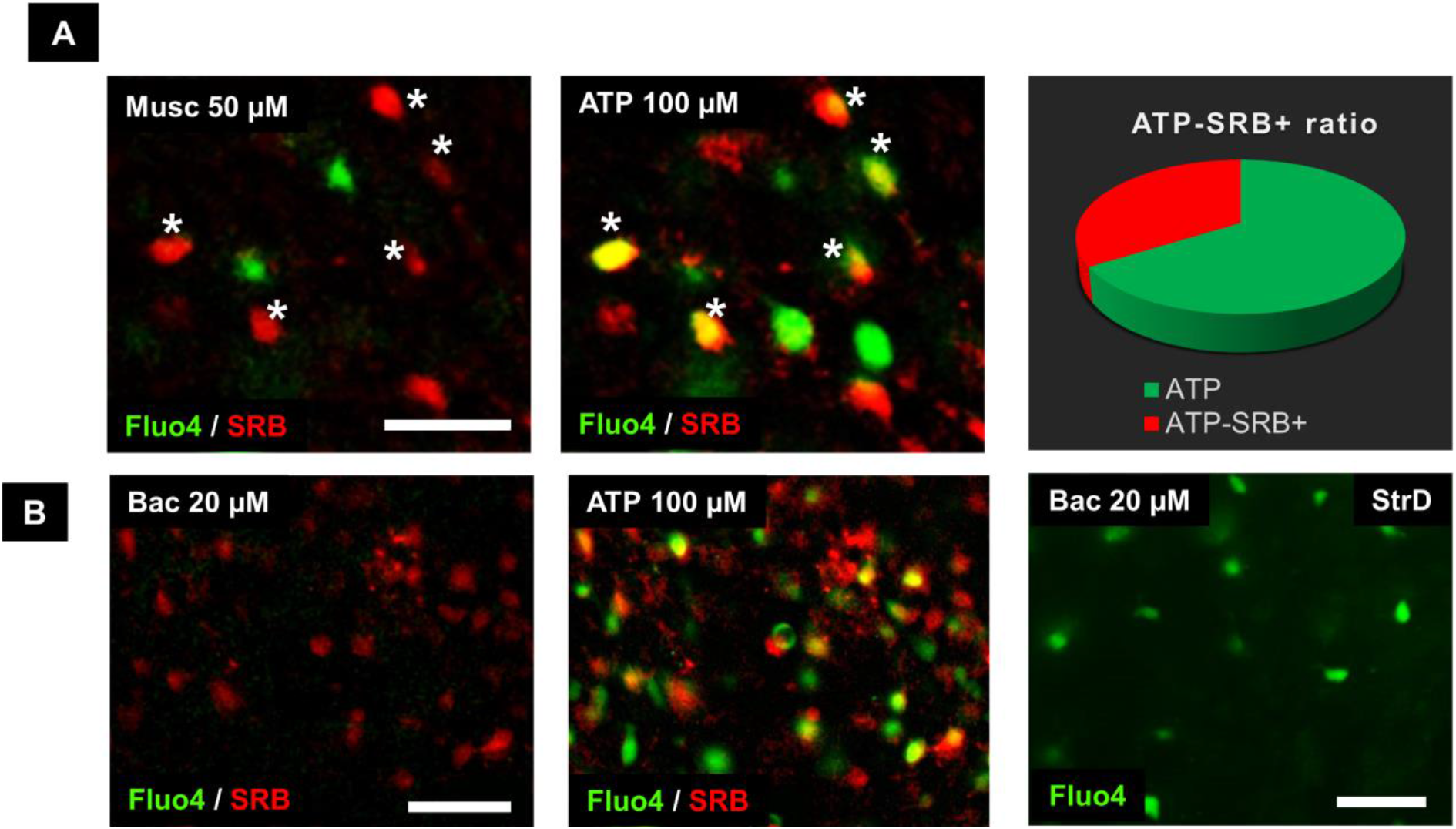
The muscimol responsive cells were not labeled by SRB. Muscimol (50 μM) responding cells were unlabeled with SRB. However, ATP (100 μM) responding cells were both SRB^+^ and SRB^−^. Asterisks indicate SRB^+^ cells (red) that did not respond to muscimol but responded to ATP (yellow). Notice that only 34% of ATP responding cells were SRB^+^ (n=6; N=5, scale bar = 20 μm). **B-** The WM cells showed no response to baclofen (20 μM). However, ATP (100 μM) activated SRB^+^ (yellow) and SRB^−^ (red) cells. As a positive control, baclofen (20 μM) responding cells were observed in the dorsal striatum (Str, P8) (n=10; N=6, scale bar = 50 μm).

### Cells from the cerebellar WM do not respond to Baclofen

Consistently, baclofen (20 μM) failed to evoke calcium responses in cerebellar WM, which contrasted to the readily evoked responses observed in SRB^+^ and SRB^−^ cells when ATP (100 μM) was added to the bath (n=8; N=4, Fig. 3B). The negative responses to baclofen in the cerebellar WM led us to test a higher concentration (100 μM), but again, cellular responses to this agonist were absent (n=6, data not shown). As a positive control, we tested baclofen on dorsal striatum and observed that cells responded to this agonist, confirming the expression of GABA_B_Rs in this brain region (n=10; N=6). Overall, we conclude that GABA_B_ receptors are not functionally expressed in cerebellar WM.

### Only NG2^+^ glia respond to the GABA_A_ agonist muscimol

The identity of GABA_A_R-expressing cells was investigated through patch-clamp whole-cell recordings performed on WM glial cells at a holding potential of −70 mV. The current profile provided initial information about cell identity and biocytin was injected at the end of the recording to test dye coupling and reveal the morphology of the cell (Fig. 4). Three cellular populations were identified. The first population corresponds to astrocytes based on GFAP-eGFP expression. These cells showed a morphology with multiple branched processes, in which biocytin diffused to neighboring cells, indicating coupling. Moreover, they showed passive linear currents with no apparent time or voltage dependence and a linear IV relation. These GFAP-eGFP^+^ cells showed no response to the GABA_A_ agonist muscimol (Fig. 4A).

**Figure 4.**
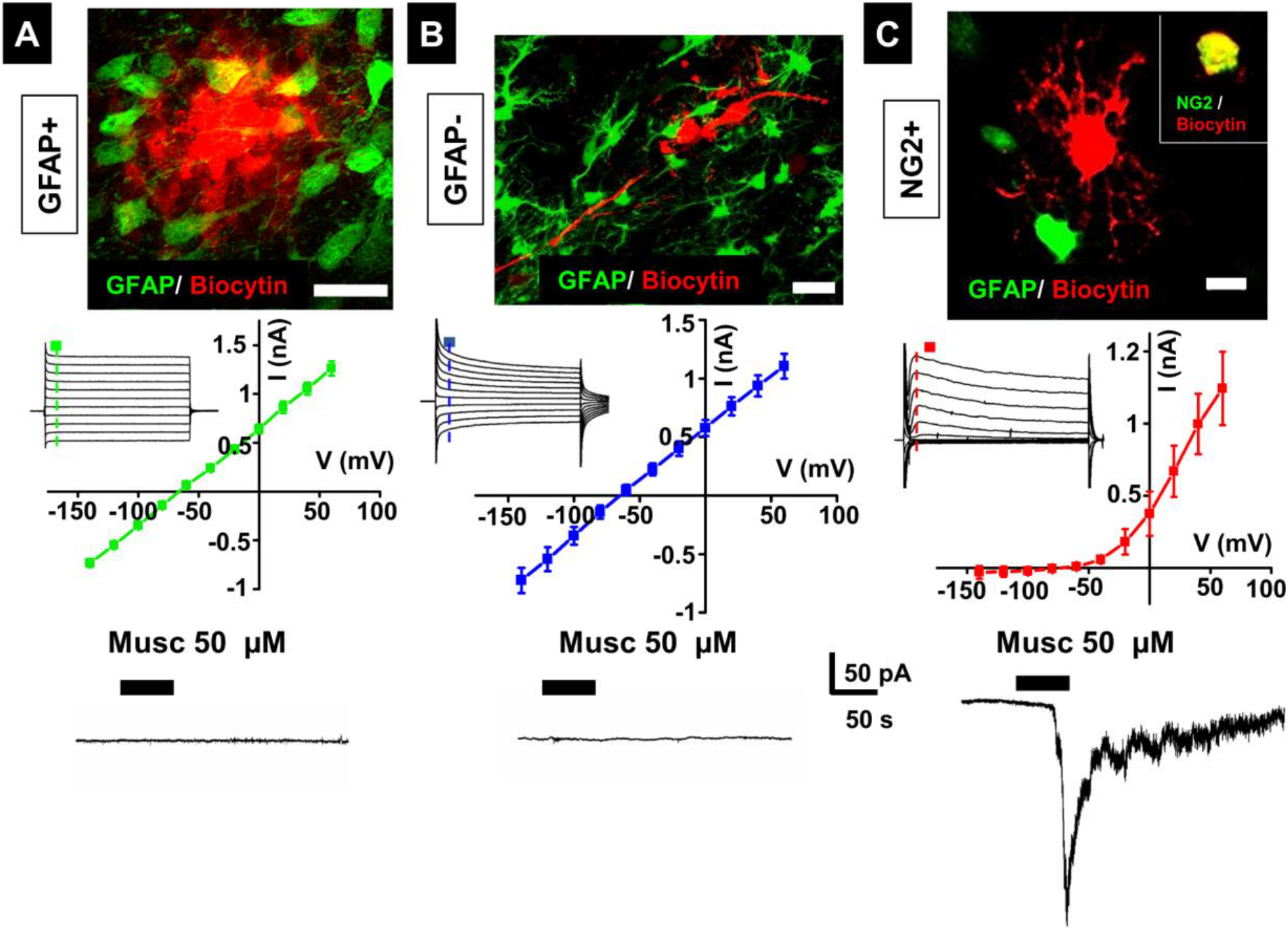
Functional expression of GABA_A_Rs is only detected in NG2 glia. **A-** GFAP^+^ cell filled with biocytin shows dye-coupling. Voltage-clamp recordings (voltage steps of 20 mV, from −140 to +60 mV) showed passive currents resulting in a linear pattern for the I-V curve, typical for astrocytes. These cells showed no response to muscimol (scale bar = 10 μm, n=9; N=8) **B-** GFAP^−^ cell filled with biocytin that diffused to a group of neighboring cells, showing coupling and revealing long horizontal processes. Voltage-clamp recordings showed currents with slow decay during voltage steps and large symmetrical tail currents that characterize oligodendrocytes. No response to muscimol was evoked (scale bar = 20 μm, n=5; N=5) **C-** NG2^+^ glia filled with biocytin showed star-shaped, highly ramified thin processes and no cell coupling (scale bar = 20 μm, n=5; N=5). Voltage-clamp recordings showed voltage-dependent currents with outward rectification in the IV curve, and muscimol evoked an inward current. Inset shows immunofluorescence of the recorded cell. The red channel corresponds to biocytin and the green channel is for NG2 antibody (scale bar = 10 μm).

The second population was GFAP^−^ in which biocytin diffused to neighboring cells that were GFAP-eGFP^−^, as they showed long parallel processes, presumably along axonal tracts, suggesting oligodendrocyte lineage. The current profile of these cells showed slow decaying inward and outward currents with no voltage dependence and a linear IV relation. Muscimol did not evoke responses in this population (Fig. 4B).

The third population was GFAP^−^ and had a star-shaped morphology with long slender processes, and biocytin did not diffuse to neighboring cells. These cells displayed voltage-dependent currents with an IV relation showing an outward rectifying current. Muscimol (100 μM) evoked inward currents (−198 ± 18 pA; n=5; N=5) indicating functional expression of GABA_A_Rs (fig. 4C). Finally, to help to determine the cell identity, biocytin-filled cells (one cell per slice) were fixed and tested for NG2 immunoreactivity. The recorded cells were immunoreactive for this marker (n=3; N=3). (Fig. 4C, inset).

### NG2 glia receives synaptic contact, but the evoked calcium wave has no effect on sPSCs frequency

NG2 glia is known to receive synaptic inputs (Muller et al 2009). Depolarization of cerebellar WM immediately evoked a postsynaptic inward current recorded in NG2 glia (50.1 +/− 4.9 pA, n=4; N=4) that was blocked with TTX (1 μM), indicating that blockage of Na channels effectively prevented neurotransmitter release from axons (Fig. 6). Additionally, spontaneous post-synaptic currents (sPSCs) were recorded from NG2 cells, these currents were fully blocked by the GABA_A_R antagonist bicuculline (100 μM. n=3; N=3; Fig 5A, B). Thus, GABA is released in the cerebellar WM and activates GABA_A_Rs expressed in NG2 glia. The axons from cerebellar WM are known to release GABA, which is consistent with our observation that showed TTX-dependent blockage of the inward currents recorded from NG2 cells but, interestingly we also observed that half of the GFAP^+^ cells express GAD67 (Fig. 1), strongly suggesting that glial cells may also release GABA within the WM. Thus, sPSCs were recorded in NG2 glia (pre-stimulus), and a calcium wave was evoked by depolarization of WM (Fig. 5C). Our results showed that the frequency of sPSCs (0.10 ± 0.05 Hz) was unchanged after the evoked calcium wave (0.18 ± 0.11 Hz; n=4 out of 11; N=4; p=0.11 Student’s t-test) (Fig. 5D).

**Figure 5.**
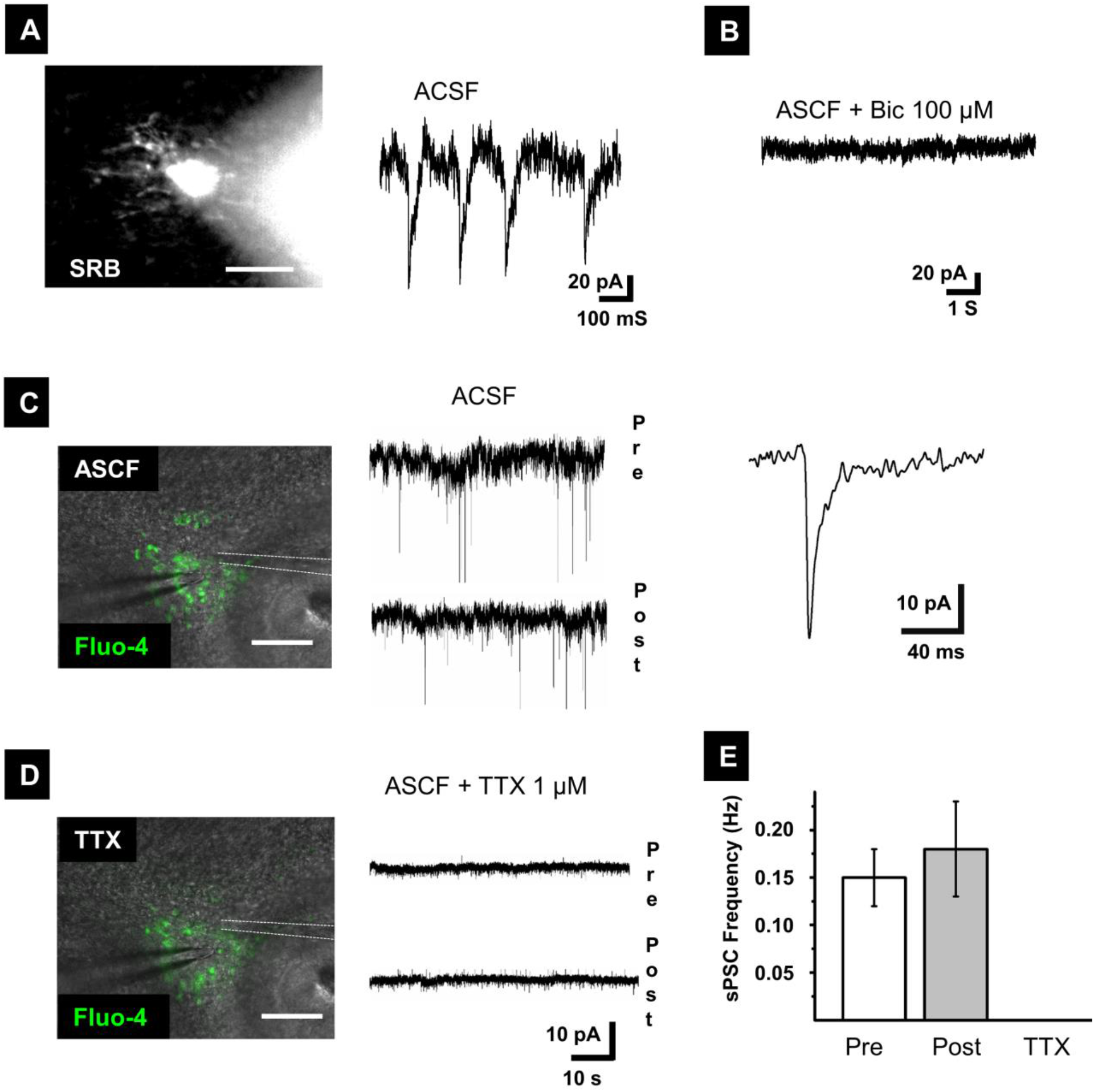
Evoked calcium waves fail to modulate sPSC frequency or signaling to NG2 glia. Representative recording of an NG2-glia showing ramified morphology revealed by SRB (scale bar = 20 μm) and sPSCs that were abolished by bicuculline (100 μM) (**B**, n=3). **C**- Representative whole-cell recording from NG2 glia showing sPSCs before (pre) and after (post) a calcium wave evoked by electrical stimulation (scale bar = 100 μm). Right, Amplification of a representative event of sPSCs. **D**- The sPSCs were abolished when TTX (1 μM) was added to the ASCF. Under these conditions, no signaling was recorded before or after the evoked calcium wave. (**E**) Summary of the experiments (n=4).

**Figure 6.**
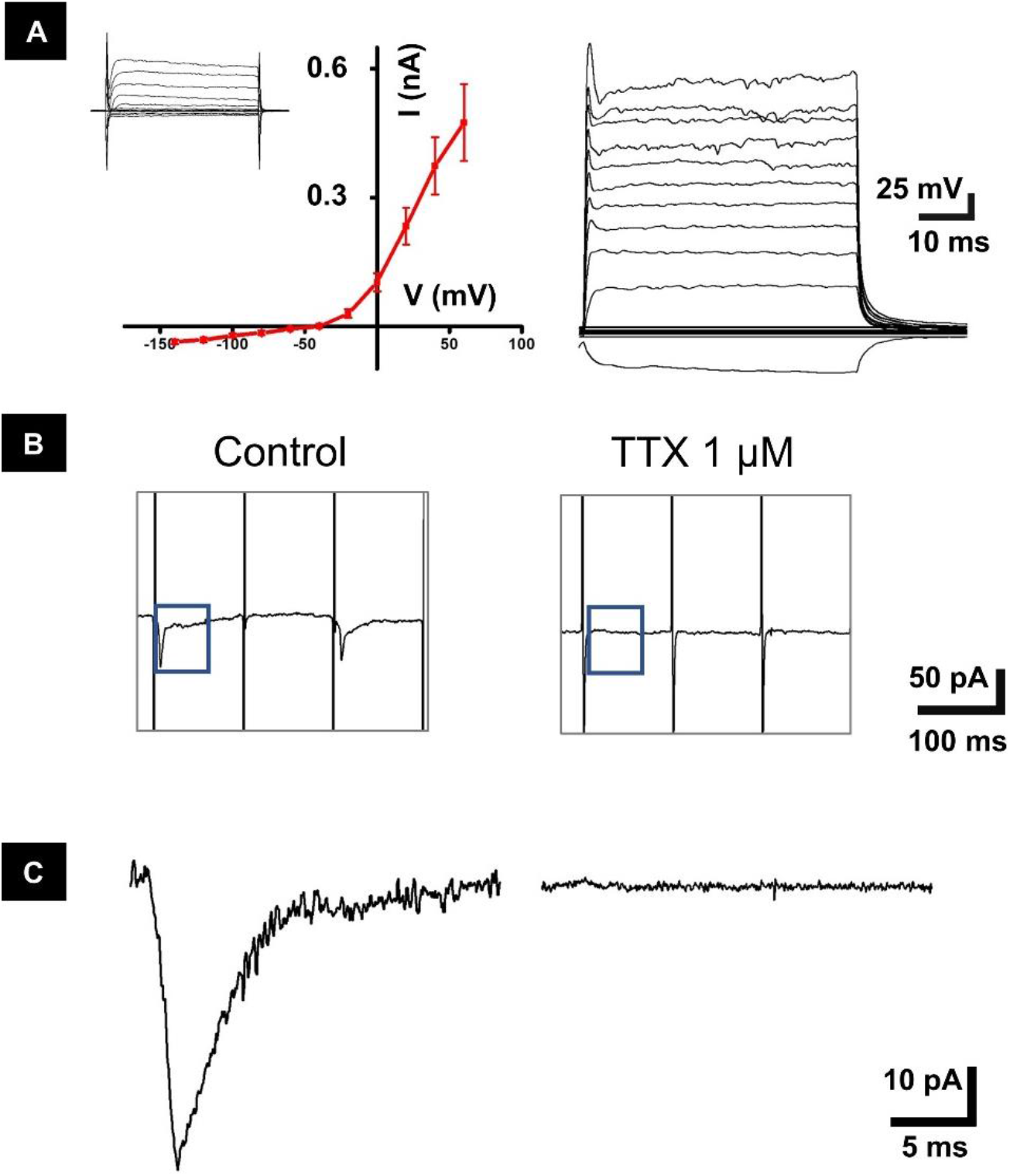
Depolarization of cerebellar WM evokes PSCs in NG2 glia. **A-** IV-curve from NG2 glia showing the typical outward rectifying current. Right: current clamp recording from the same cell. Current injection failed to induce action potentials (steps from −80 to 400 pA, 30 pA/step) **B**- Evoked PSCs in NG2 glia after depolarization of cerebellar WM (200 μA, 10 Hz, 2 s). The evoked PSCs were abolished when TTX (1 μM) was added to the ACSF (n=4; N=4). **C**- Amplification of the evoked PSC and the recording in ACSF+TTX.

Our next step was to block the axonal mediated response with TTX (1 μM) while the calcium wave was kept promoting gliotransmitter release. However, TTX completely abolished sPSCs from NG2 glia in which the calcium waves did not suffice to elicit detectable currents (n=5; N=5; fig. 5D, E).

## Discussion

In this study, we investigated the GABAergic signaling of glial cells from cerebellar WM. Our first result showed that GAD67 is densely expressed in early postnatal development (P7-9), confirming previous observations in GAD67-GFP knock-in mice (Yamanaka et al., 2004) that showed abundant GFP-positive cells in the WM at P5-15 that gradually reduced and were scarce by P21. Interestingly, we also found that half of the GFAP^+^ cells express GAD67, suggesting that GABA is synthesized within the WM and possibly released from this group of cells. This set the basis to investigate the functional expression of GABA receptors in glial cells of cerebellar WM. Calcium imaging and selective pharmacological activation revealed responses to muscimol but not to baclofen, indicating that GABA-signaling in this area is mainly mediated by GABA_A_Rs but not GABA_B_Rs. This contrasts to previous studies reporting the expression of GABA_B_Rs in glial cells from other brain regions. For example, a population of cortical astrocytes expresses these receptors in slices from P15-20 and P30-60 mice (Mariotti et al., 2016). Likewise, the GABA_B_ agonist baclofen elicited calcium transients in hippocampal astrocytes (Perea et al., 2016), oligodendroglial lineage cells (O4^+^) from the periventricular WM (Luyt et al., 2007) and striatal astrocytes (Nagai et al., 2019). In this study, we confirmed that baclofen generates a robust response in the striatum, but fails to elicit cellular responses in cerebellar WM. Altogether these results strongly suggest that functional expression of GABA_B_Rs is absent in cerebellar WM, at least in early postnatal development.

### GABA_A_ is functionally expressed in NG2 glia

We tested the functional expression of GABA_A_Rs in astrocytes, oligodendrocytes and NG2 glia. Microglia was discarded in this study because there is no evidence of functional GABA_A_R expression in these cells (Cheung et al., 2009; Kettenmann et al., 2011). On the other hand, several reports have shown GABA_A_-mediated responses in astrocytes (Fraser et al., 1994, Müller et al., 1994, Bolteus and Bordey, 2004, Reyes-Haro et al., 2013a), oligodendrocytes (Berger et al., 1992, Arellano et al., 2016, Habermacher et al., 2019) or NG2 glia (Luyt et al., 2007, Balia et al., 2013, Zonouzi et al., 2015, Habermacher et al., 2019). However, little is known about the functional expression of GABA_A_Rs in cerebellar WM (Zonouzi et al., 2015).

Glial cells dominate the cellular composition of WM in which they provide 99% of the cell somas (Sturrock, 1976; Reyes-Haro et al., 2013). In our studies calcium imaging showed that muscimol elicited calcium transients in cerebellar WM, indicating functional expression of GABA_A_Rs. However, this experimental set did not provide information about the identity of the responding cells.

Our next step was to stain astrocytes *in situ* with SRB. This methodology (Appaix et al., 2012) has demonstrated to be reliable but with some limitations, since it can also mark oligodendrocytes (Hill & Grutzendler, 2014, Hülsmann et al., 2017). We observed that SRB^+^ cells did not respond to muscimol, suggesting that astrocytes lack functional expression of GABA_A_Rs.

Next, patch-clamp recordings were performed on astrocytes, NG2 glia, and oligodendrocytes. This technique provides information about the current profile, morphology and dye coupling. Additionally, we tested the functional expression of GABA_A_Rs and only found muscimol-mediated responses in NG2^+^ glia, indicating that astrocytes and oligodendrocytes lack functional expression of this receptor. This study is the first one to test the functional expression of GABA receptors in three different types of WM glial cells and reveals specific GABA_A_-mediated signaling for NG2 glia only. This is consistent with our calcium imaging studies showing that SRB^+^ cells did not respond to muscimol and was corroborated with whole-cell patch-clamp studies in astrocytes and oligodendrocytes.

The lack of muscimol response in astrocytes was surprising considering that functional expression of GABA_A_Rs was demonstrated in Bergmann glia at P5-P7 and GABA_A_- mediated signaling at P20-P30 was drastically reduced or undetectable (Müller et al., 1994), suggesting that while GABA_A_ signaling is important through early postnatal development of the cerebellum, there is an apparently different regulation for WM. In support of this hypothesis, functional expression of GABA_A_Rs was also reported in ependymal glial cells from the cerebellum located at the roof of the fourth ventricle (Reyes-Haro et al., 2013a), while immunofluorescence studies revealed the expression of the GABA_A_ρ1 subunit in astrocytes from the granular layer of the cerebellar lobule X (Pétriz, et al., 2014). Astrocytes from WM give rise to interneurons of the molecular layer during the postnatal development of the cerebellum (Silbereis et al., 2009). In contrast, migration tracking studies in organotypic cultures of hippocampal slices showed that astrocytes with neurogenic activity regulate migratory direction and speed through GABA_A_Rs (Bolteus and Bordey, 2004). Our results indicate that WM astrocytes do not respond to GABA, this suggests that a different mechanism must operate for controlling the migration of GFAP^+^ neuronal precursors in the cerebellum.

Another interesting observation was the coupling between GFAP^+^ and GFAP^−^ cells revealed by diffusion of biocytin. We consider two possible explanations for this result. First, we know that not all the astrocytes are labeled by the eGFP reporter in the transgenic model used in our study (Nolte et al., 2001). Second, astrocyte-oligodendrocyte coupling via gap junctions was observed in the corpus callosum and cerebellar WM (Maglione et al., 2010, Tress et al., 2012). Our results suggest to not discard either astrocyte GFAP^+^-astrocyte GFAP^−^ nor astrocyte-oligodendrocyte coupling, adding a new layer of complexity between different glial cells.

### NG2 glia from cerebellar WM receive synaptic input

Neuron-NG2 glia communication in the cerebellum was first described in the molecular layer, where depolarization of climbing fibers evoked glutamatergic postsynaptic currents (Lin et al., 2005). Later, GABAergic synaptic input was recorded in NG2 glia from cerebellar WM (Zonouzi et al., 2015). We confirmed this result and found that GABA_A_R-mediated signaling is selective for NG2 glia in cerebellar WM. This was a surprising result considering that functional expression of GABA_A_Rs was reported in glioblasts and oligodendrocytes from the corpus callosum, another WM tract (Berger et al., 1994). Accordingly, neuron-oligodendrocyte co-cultures showed functional expression of GABA_A_Rs, although isolation of oligodendrocytes resulted in a loss of GABA-mediated responses (Arellano et al., 2016), indicating that neuronal contact is necessary for functional expression of GABA_A_Rs. Moreover, proliferation and myelination of oligodendroglia is regulated by GABA_A_Rs in cortical cultures (Zonouzi et al., 2015; Hamilton et al., 2017). However, different actions were reported for *in vitro* vs *in vivo* studies when a GABA_A_R antagonist was used. For example, the application of gabazine increased the number of CC1^+^ oligodendrocytes and myelination in cortical cultures (Hamilton et al., 2017), while *in vivo* studies showed that bicuculline increased NG2 glia and decreased CC1^+^ oligodendrocytes in the cerebellum (Zonouzi et al., 2015). These differences may be explained by regional particularities or differences between *in vitro* and *in vivo* studies.

### Evoked calcium waves fail to signal NG2 glia

Glial cells exhibit a form of long-distance signaling in which propagation of calcium transients occurs among cells. These calcium waves were described in cortical WM and propagate via gliotransmitter release (Schipke et al., 2002; Haas et al., 2006). As shown in our study, half of the cerebellar WM astrocytes express GAD67, which may lead to local synthesis of GABA. GABA signaling from glial cells is carried out by vesicular release, reverse activity of GABA transporters or release of GABA through the bestrophin-1 channel (Song et al., 2013; Lee et al., 2010; Yoon et al., 2011). We ruled out glia-mediated GABA transmission since no responses were detected in NG2 cells when TTX was added to the aCSF, which indicated that GABAergic transmission into NG2 glia is mainly neuronal and that glial calcium waves do not elicit any measurable electrophysiological response. Our results agree with a previous report suggesting that NG2 glia from cerebellar WM receives synaptic input from local GAD65+ interneurons (Zonouzi et al., 2015). Interestingly, we only recorded sPSCs in four out of eleven NG2 cells. This proportion is slightly lower than that of a previous study where 20 out of 31 Ds-Red-NG2+ cells showed postsynaptic activity (Zonouzi et al., 2015). This signaling may reflect a specific regulatory pathway for controlling NG2 cell development. The action of GABA_A_Rs in NG2-glia may be time restricted to early postnatal ages, as shown in other brain regions (Orduz et al., 2015, Parolisi and Boda, 2018). NG2 glia gives rise to oligodendrocyte precursor cells which keep proliferating and differentiating into myelinating oligodendrocytes even in the adult life. In early postnatal development of the cerebellum, we have shown communication between neurons and NG2s that receive GABAergic synaptic input, thus providing a probe of the relevance of this signaling in neuron-glia communication.

In sum, three glial cell types were tested for functional expression of GABA receptors. Only NG2 glia detects GABA signaling through GABA_A_Rs because neither astrocytes nor oligodendrocytes responded to this neurotransmitter. Functional expression of GABA_B_Rs is absent in cerebellar WM and GAD67 is expressed in half of the astrocytes. Axon-NG2 glia signaling is mediated through GABA_A_Rs, but glia-glia signaling was not detected.

## Acknowledgments

Francisco Emmanuel Labrada-Moncada is a doctoral student from Programa de Doctorado en Ciencias Biomédicas, Universidad Nacional Autónoma de México (UNAM) and received fellowship from CONACYT (640190). We are grateful to AE. Espino, A. Castilla León, N. Hernández Ríos, M García Servín, L. Casanova Rico and M.L. Lara Ayala for their technical support. We are indebted to Dr. R. Arellano Ostoa for allowing us to use his histological facilities. The authors thank LCC Jessica González Norris for proofreading the English version of this manuscript. This work was supported by PAPIIT-UNAM to AMT (IN200913) and DRH (IN201915 and IN205718).

